# Multi-modal characterization and simulation of human epileptic circuitry

**DOI:** 10.1101/2020.04.24.060178

**Authors:** Anatoly Buchin, Rebecca de Frates, Anirban Nandi, Rusty Mann, Peter Chong, Lindsay Ng, Jeremy Miller, Rebecca Hodge, Brian Kalmbach, Soumita Bose, Ueli Rutishauser, Stephen McConoughey, Ed Lein, Jim Berg, Staci Sorensen, Ryder Gwinn, Christof Koch, Jonathan Ting, Costas A. Anastassiou

## Abstract

Temporal lobe epilepsy is the fourth most common neurological disorder with about 40% of patients not responding to pharmacological treatment. Increased cellular loss in the hippocampus is linked to disease severity and pathological phenotypes such as heightened seizure propensity. While the hippocampus is the target of therapeutic interventions such as temporal lobe resection, the impact of the disease at the cellular level remains unclear in humans. Here we show that properties of hippocampal granule cells change with disease progression as measured in living, resected hippocampal tissue excised from epilepsy patients. We show that granule cells increase excitability and shorten response latency while also enlarging in cellular volume, surface area and spine density. Single-cell RNA sequencing combined with simulations ascribe the observed electrophysiological changes to gradual modification in three key ion channel conductances: BK, Cav2.2 and Kir2.1. In a bio-realistic computational network model, we show that the changes related to disease progression bring the circuit into a more excitable state. In turn, we observe that by reversing these changes in the three key conductances produces a less excitable, “early disease-like” state. These results provide mechanistic understanding of epilepsy in humans and will inform future therapies such as viral gene delivery to reverse the course of the disorder.

---

Epilepsy is one of the most common neurologic ailments and temporal lobe epilepsy (TLE) is its most commonly diagnosed form affecting approximately 65 million people worldwide. Despite considerable advances in the diagnosis and treatment of such seizure disorders, the cellular and molecular mechanisms leading to TLE-related seizures remain unclear. Notably, approximately 40% of TLE patients exhibit pharmaco-resistance, i.e. lack of response to conventional anticonvulsive treatments (*1*). Hippocampal sclerosis (HS), a neuropathological condition associated with cell loss and gliosis, has been linked to increased occurrence of TLE with elevated degree of HS constituting a hallmark of disease progression. The sclerotic hippocampus is thought to be the most likely origin of chronic seizures in TLE patients and is the target of temporal lobe resection intervention. The efficacy of these surgical interventions (i.e., 65-80% of TLE patients become seizure-free) further implicates the sclerotic hippocampus as a prominent member of the pathophysiological network (*2, 3*). Patients with increased HS typically suffer from more frequent seizure activity (*4–8*), an observation also replicated in rodents (*9*). Furthermore, HS has been associated with a spectrum of adverse effects such as longer epilepsy duration (*10–13*), an earlier stage of onset (*8, 14, 15*) as well as the presence of early aberrant neurological insults such as febrile convulsions (*8, 14, 15*).

To mechanistically understand the link between disease progression and increased seizure propensity in TLE patients, we studied live human brain tissue excised during temporal lobectomy (Fig. 1 and Table S1). The excised brain tissue was quantified neuropathologically via the Wyler grade or WG (WG range: 1-4) (*16*) based on light microscopic examination of the mesial temporal damage in temporal lobectomy specimens (Fig. 1A): degree 1 (WG1) corresponds to none/mild HS, while degree 3 or 4 (WG4) corresponds to severe HS. We use a data generation and analysis platform (*17*) to characterize disease-related features in gene expression (RNA-seq), electrophysiology, morphology, spine density and gene expression at single-cell resolution. Specifically, we concentrated on granule cells (GCs) of the dentate gyrus (DG), a hippocampal region. Studies in animal models and human neurosurgical specimens have implicated DG GCs in the generation and support of seizure activity by a plethora of mechanisms spanning from altered excitability, changes in morphology, protein expression, neuronal loss, synaptic re-organization and altered connectivity patterns (*18–20*). We pursued the multi-modal characterization of human GCs across patients and report how their properties change with disease progression, i.e. increasing WG (Fig. 1B and 1C, S1 and S2).

**Figure 1:**
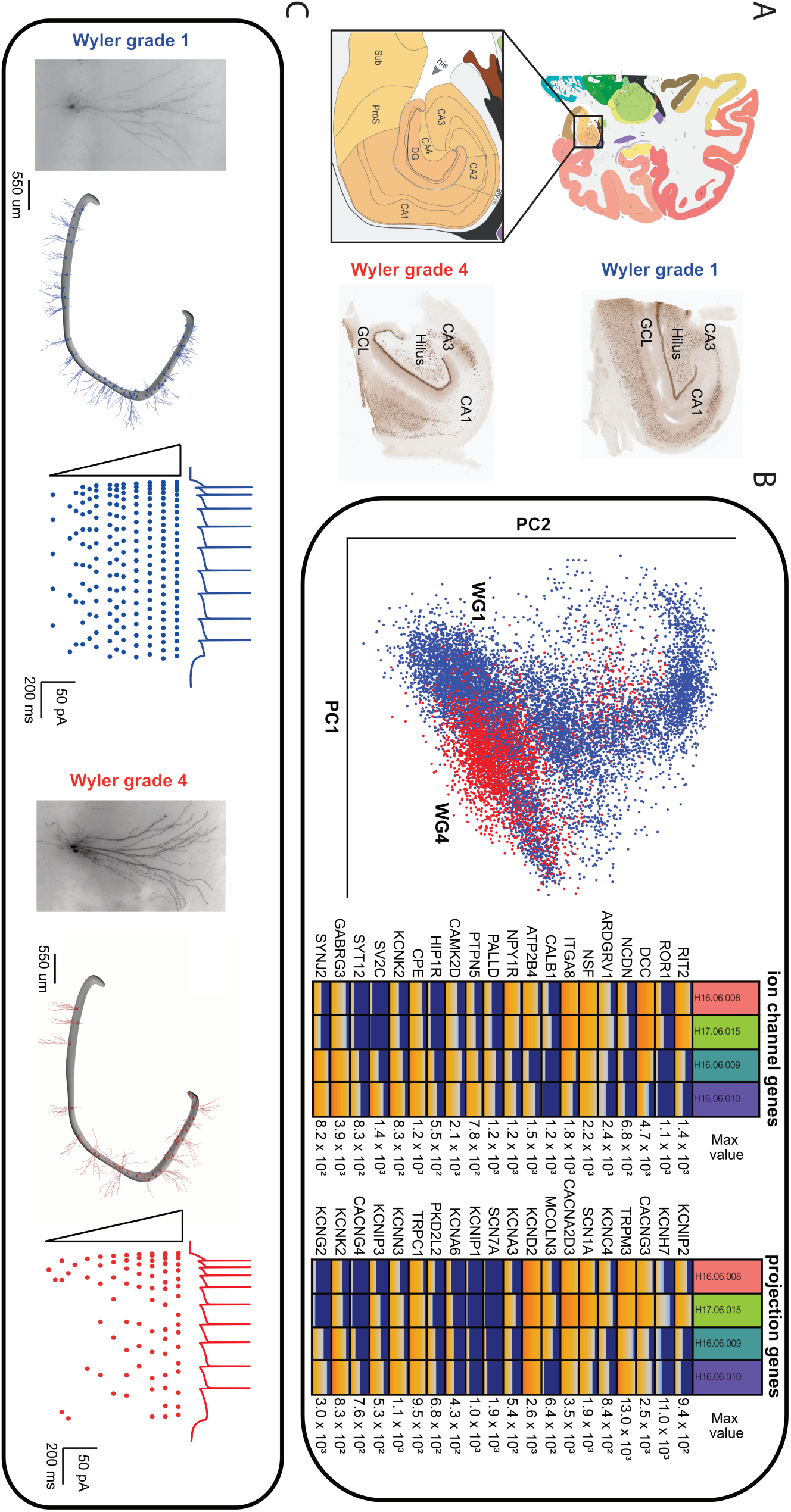
Human data collection from drug-resistant temporal lobe epilepsy patients. (A) Human dentate gyrus (DG), Cornu Ammonis (CA) 1-4, pro-subiculum (ProS) and subiculum (Sub) areas typically resected during temporal lobectomy. (B) Single-cell gene expression data of 230 ion channel-coding and 1600 projection-coding genes specific for human granule cells separate cells with respect to disease state. Left: principle component analysis of the subset of 1600 projection genes for nuclei with the expression profiles mapped to human DG granule cells (n_cells_(WG1) =8760, n_cells_(WG4)=3251; see Supplementary Material). Right: expression level of ion channel and projection genes for different patients (n=2 for Wyler grade 1, n=2 for Wyler grade 4). Rows correspond to genes and columns to patients. Expression levels are shown in counts per million (CPM) base pairs using fire plot. (C) Human hippocampal slices retrieved from surgical specimens (DG GC layer designated by neuronal nuclei stain). Hippocampal sclerosis (HS), as assessed by Wyler grade is associated with cell loss in hippocampal subfields such as CA1 (e.g., compare cellular density in area CA1 for WG1 vs. WG4 in panel A). Locations along the human DG where whole-cell patch-clamping and morphological reconstructions were gathered across patients. Somatic trace (top) and raster (bottom; lines: experiment trials from the same cell) from whole-cell patch-clamp experiments involving intracellular somatic current dc injection of increasing magnitude in DG granule cells from a WG1 (blue) and a WG4 (red) patient case.

## Single-cell gene expression changes in human granule cells with TLE progression

To characterize the expression profile in the human epileptic hippocampus, we analyzed the gene expression data of 230 ion channel-coding and 1600 projection-coding specific for GCs (Fig. 1B). Analysis of the sc RNA-seq data set exhibits that a large number of genes are differentially expressed with disease progression in GCs clearly separating WG1 and WG4 (Fig. 1B). In addition, we found that the proportion of GCs was larger in WG1 compared to WG4 patients (57% vs. 30% of all cells, respectively). These findings indicate that a number of GC properties change with disease progression as well as the progressive cells loss in human hippocampus with degree of hippocampal sclerosis. To study the cellular changes implicated by the sc RNA-seq data, we performed a sc electrophysiology and morphology survey of GCs in the same human brain slices and examined how these properties change with disease progression (Fig. 1C).

## Excitability changes in granule cells with TLE progression

We employed whole-cell patch-clamping in human DG GCs imposing a battery of intracellular stimuli to assess electrophysiological changes associated with HS. The standard protocol (1 s-long dc current injections of different amplitudes) was applied to 112 patch-clamped GCs (Fig. 2A). A set of 31 electrophysiological features was extracted for every experiment (Table S2) and evaluated over the range of injected current amplitudes. Based on these electrophysiology features, we observed robust separation between WG1 and WG4 cases (Fig. 2B-C; tSNE: t-distributed stochastic neighbor embedding (*21*)). To identify the electrophysiology features separating WG1 vs. WG4, we used two methods: i) pairwise feature comparison in combination with Mann-Whitney U-testing (*p*<0.05/*n*, Bonferroni-corrected for multiple comparisons *n*), which resulted in 9 out of 31 features exhibiting statistically significant difference (Fig. 2D); ii) random forest classification reaching classification accuracy of 81% (chance level: 50%; classifier trained on all electrophysiological features; out-of-bag error; Fig. 2E). When comparing the classifying features resulting from the two methods, we found that 5 out of 9 statistically significant features were shared (Fig. 2D and 2E). The most prominent electrophysiological features affected by disease progression are the spike frequency-current (“f-I”) gain (WG1 GCs have a higher gain than WG4) and the time-to-spike (WG4 GCs have a shorter time-to-spike duration than WG1), two prominent excitability parameters of a neuron (Fig. 2F). To cross-validate against patient-specific effects, patient-out-validation (classifier trained on data from 6 patients predicting WG-score of 7^th^ patient) classification performance reached 71% (chance level: 50%; Table S3).

**Figure 2:**
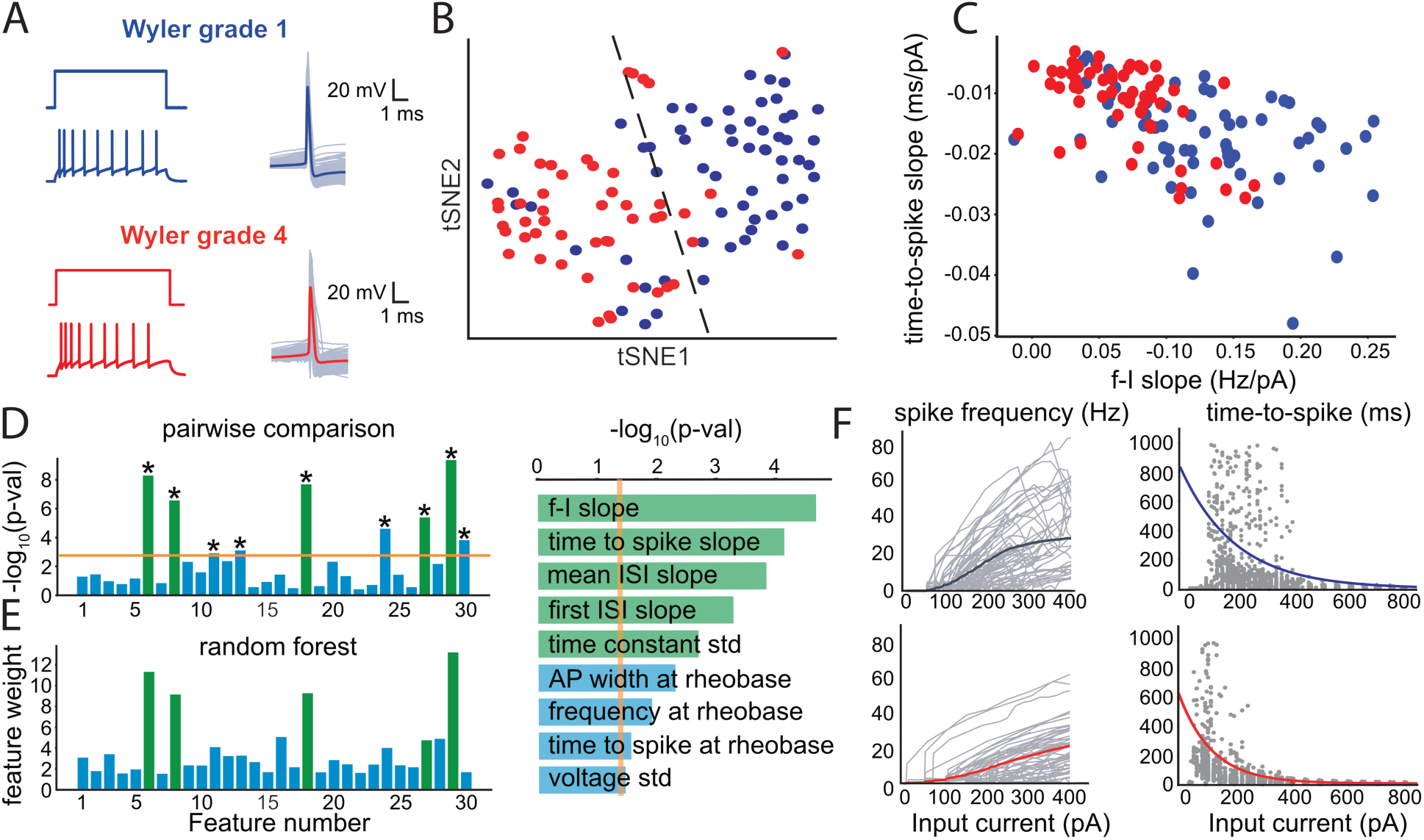
Electrical behavior of human granule cells change with disease severity and degree of histological degeneration. (A) Whole-cell patch-clamping of DG GCs across patients and somatic spiking responses to 1s-long dc current injections (blue: WG1; red: WG4). Intracellular somatic spike waveform (thick line: average waveform; thin lines: individual waveforms from a particular experiment). (B) tSNE visualization using 30 electrophysiological features from 112 human DG GCs (WG1: 61; WG4: 51 cells; blue: WG1, red: WG4; broken line: *k*-means decision boundary in tSNE space between the two main clusters; Table S2). (C) Two key electrophysiology features leading to WG-separation: time-to-spike slope (ms/pA) vs. spike-frequency slope (Hz/pA). (D) Pairwise comparison of electrophysiological features between WG1 and WG4 GCs across feature space (left) and in decreasing level of statistical significance (right). Statistical significance is calculated using Mann-Whitney U-test with the significance level (orange line) Bonferroni-corrected for the number of features (* designates *p*-value < 0.05/30=0.002). (E) Electrophysiology feature weights of random forest classifier. Shared (green) and non-shared (blue) electrophysiology features between pairwise comparison and random forest classifier among the 10 most influential ones are shown (one-out-of-bag validation; classification accuracy = 81%) (F) Spike-frequency and time-to-first-spike response to dc current injections for WG1 (top) vs. WG4 (bottom) exhibit altered GC-excitability (thick line: average; thin lines: mean response curves from individual experiments; grey points: first spike time in all experimental sweeps).

## Altered granule cell morphology and spine density

Next, we ask to what extent the morphology of human GCs is affected by disease progression. To do so, we reconstructed the dendritic morphology of 102 single GCs (Fig. 3A) and used a set of 49 morphology features to assess WG-dependent alterations (Table S4). We found that morphologies differ between neurons from different levels of disease progression (Fig. 3B-C). Following the same methodology as for electrophysiology features, we used two tests to detect the features leading to the morphology separation (Fig. 3D-E). Random forest classification (out-of-bag error) resulted in morphology-based classification performance of 74%, i.e. comparable to that of electrophysiology features. Notably, most features point to disease progression correlating with thicker (i.e. increased surface area and volume) GC morphologies. When training on both electrophysiological (Fig. 2) and morphology features (Fig. 3) for the subset of GCs (*n*=77) where both data modalities are present, classifier performance reached 81% (Fig. S3).

**Figure 3:**
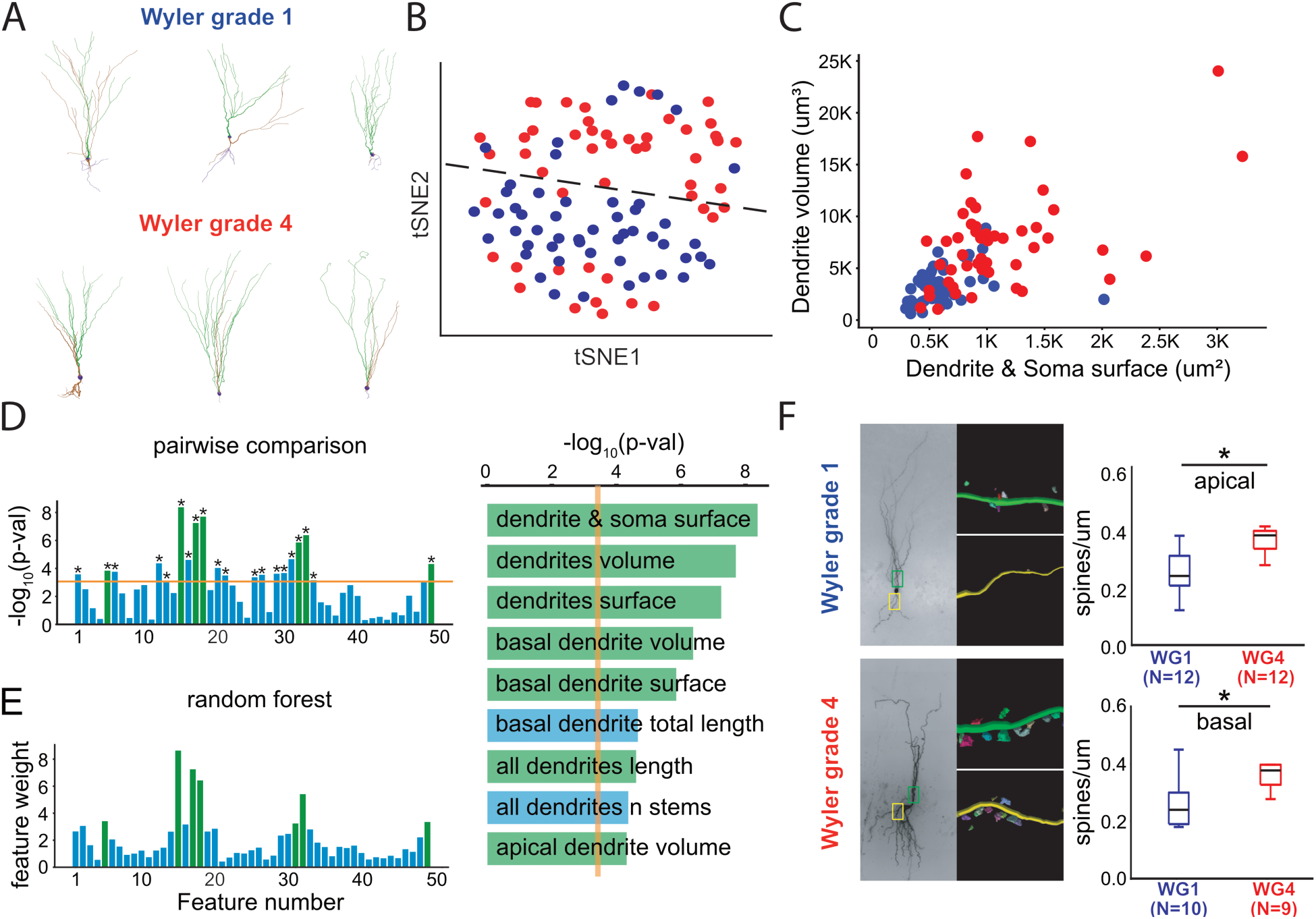
Morphology and spine density of human granule cells are affected by disease severity. (A) Reconstructed morphologies of GCs from mild and severe HS donors (green: apical dendrite region; brown: basal dendrite region; violet: axonal region). (B) tSNE visualization based on 99 single-cell morphological dendritic features of a total of 102 GC reconstructions (see Table S3) illustrates separation (blue: WG1, 52 reconstructions; red: WG4, 50 reconstructions; broken line: *k*-means clustering). (C) Two morphology features leading to WG-separation: dendritic volume (μm^3^) vs. dendritic and soma surface area (μm^2^). (D) Pairwise comparison between WG1 and WG4 morphologies (two-sample Mann-Whitney U-test, orange line: significance level Bonferroni-corrected for the number of features, *p*-value < 0.05/49=0.001). (E) Weights of the random forest classifier trained solely on morphology features for WG1 and WG4 granule cells, out-of-bag classification accuracy 74 % (right: features shared between pairwise comparison and random forest classifier shown in green, non-shared ones in blue). (F) Spine density along human granule cells depends on WG. (Left) Morphological reconstructions of single neurons including region-dependent spine density estimation (yellow box: basal region 100 μm after GC layer; green box: apical region 100 μm after GC layer). Individual reconstructed sections with spines along various regions for two WG1 and WG4 neurons are also shown. (Right) Spine density along apical and basal dendrites (box plots: median, first and third quartile and data max/min; significance level = 0.05; *p*-value calculated via Mann-Whitney U-test, * designates *p* < 0.05).

To what extent is synaptic input along GCs affected by disease progression? To determine the amount of excitatory synaptic input along GCs we estimate the spine density in a subset of the morphological biocytin reconstructions (Fig. 3F). Average spine density increases approximately two-fold from WG1 to WG4. This suggests an increase in synaptic drive of GCs with disease progression. Notably, differences in spine density occurred both along the apical and basal dendrites. Overall, our results from spine density measurements indicate that DG GCs in severe stages of HS receive increased excitatory synaptic drive compared to mild HS.

## Ion conductance changes

To investigate the functional consequences of TLE and HS on electrophysiological, morphological and connectivity properties of human GCs, we generated a series of biophysically detailed, anatomically realistic, conductance-based single-neuron models (*22*). To produce faithful single-neuron models, a key question is which ionic conductances to account for. Our starting point for the ionic conductance recipe relied on a detailed GC model based on the rodent literature that involves 7 voltage-dependent K-channels, 4 voltage-dependent Ca-channels, 2 Ca-dependent K-channels as well as two voltage-dependent Na- and Na/K (HCN) channels distributed along different parts of the morphology (Fig. 4A, (*23*)). To assess to what extent these conductances are truly present in human GCs, in a second step, we analyzed fluorescence activated cell sorted single-cell RNA-seq data from four human brain specimens (*24*) to test for the expression of genes associated to the ionic conductances used in our models. We found that the vast majority of the genes associated with the conductances are indeed expressed in human DG neurons (Fig. 4B).

**Figure 4:**
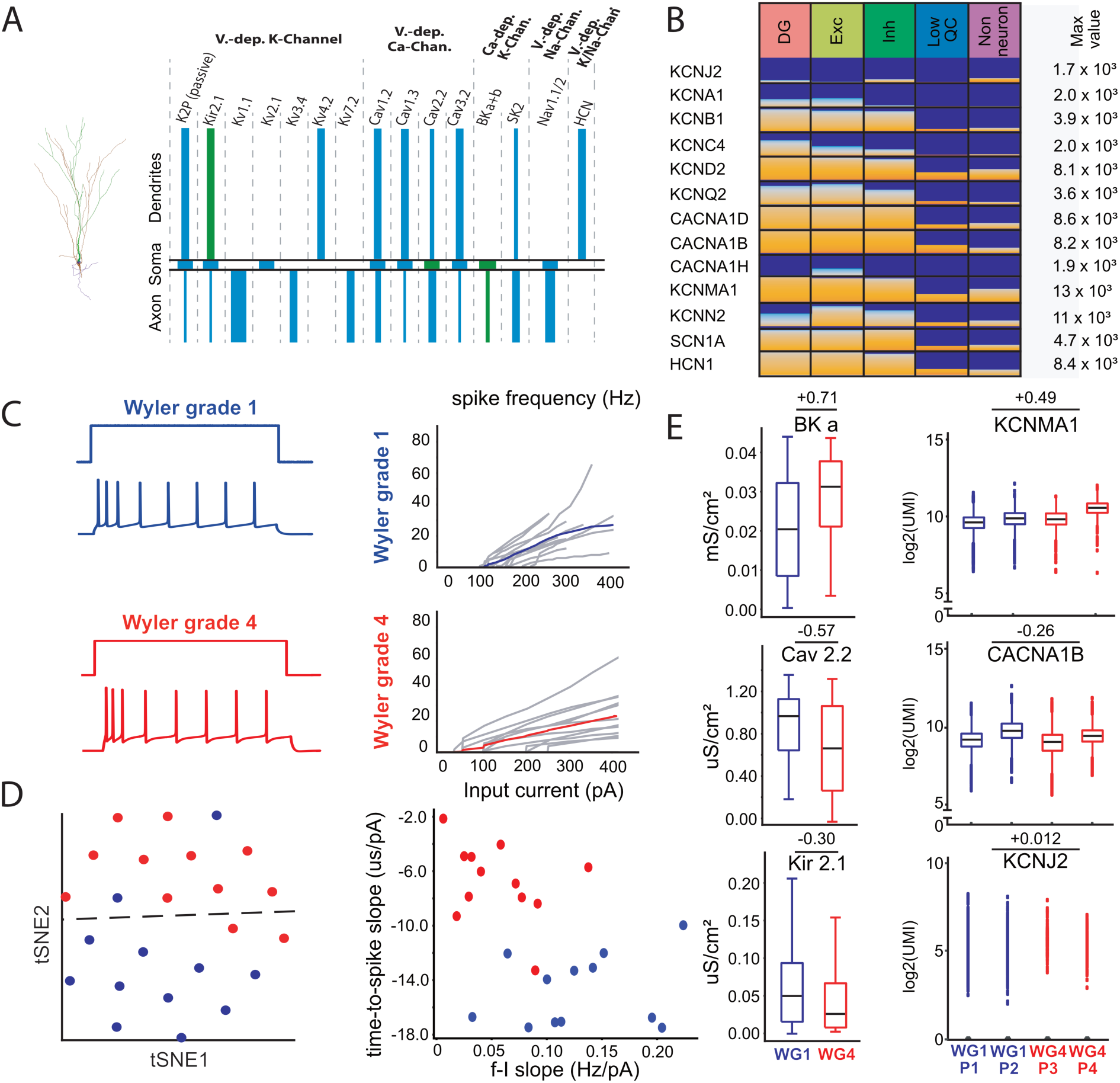
Computational modeling and single-cell RNA-sequencing implicate alterations in specific conductances with disease progression. (A) Single-cell model setup using 15 ionic conductances distributed along axonal, somatic and dendritic sections of the morphology reconstruction. A genetic optimization framework was used to estimate conductance properties (Fig. S4) and set parameter values (Table S5) at the single-cell level. Colors indicate ion conductances responsible for model-based WG-separation based on conductance values (green: statistically significant differences between WG1 vs. WG4 conductances; blue: no statistical significance; *p*-value calculated using Mann-Whitney U-test; significance level: *p* < 0.05/61=0.001, Bonferroni-corrected for multiple comparisons). (B) Expression profiles of nuclei from human DG mapped to major cell classes for a subset of genes associated with the ion channels from the GC model setup (panel A). Columns correspond to different cell classes (left-to-right): all cells in human DG, excitatory and inhibitory neurons and non-neuronal cells. Expression levels are shown in counts per million (CPM) base pairs using fire plot. (C) Simulated electrophysiological recordings from the GC models resulting from the 3-stage model generation workflow (Materials and Methods). (Left) GC model responses to 1 s-long dc current injection. (Right) f-I curves from the GC models (thick line: mean; grey lines: individual responses). (D) tSNE visualization (left) of the electrophysiology features from the GC models based on their response to 1 s-long somatic dc current injections (blue circles: WG1, red: WG4; left, broken line: *k*-means clustering). The two most prominent electrophysiology features (right) separating GC models based on WG are the same ones as for experiments (Fig. 2, C and D) (**E**) Pairwise comparison of differentially expressed conductances as predicted by the computational models between WG1 and WG4 and 10X single nucleus RNA-seq data for BK (KCNMA1), Cav2.2 (CACNA1B) and Kir2.1 (KCNJ2). (Left) For model-based ionic conductance comparisons, 10 best GC models per cell were chosen (10 models per cell * 12 cells per WG = 120 models per WG; Materials and Methods). (Right) Expression profiles of human GCs measured via single-cell RNA-seq from 2 WG1 (blue) and 2 WG4 (red) patients. For BK, Cav2.2, and Kir2.1 (log2-fold change shown; boxes: 25^th^/75^th^ percentile; whiskers: +/− 1.5x interquartile range).

To generate human GC single-neuron models we used a 3-stage approach leveraging a genetic optimization framework which relies on comparisons between experimental and model electrophysiology features for a particular morphology reconstruction (Fig. S4, (*25–27*)). Using this optimization framework, we generated GC models accounting for active ionic conductances along their entire soma, axonal and dendritic arbor. In total, we developed 12 single-cell models per WG (data originating from multiple patients) in an unbiased manner that captured a multitude of experimental observables such as differences between WG1 vs. WG4 in terms of spike frequency response to increasing injection currents, time-to-spike, etc. (Fig. 4C and Fig. S5) As a result, when classifying electrophysiology features from these models, WG1 vs. WG4 models are clearly separated and, importantly, the features leading to this separation are in agreement with the ones leading to separation of experimental electrophysiology features (Fig. 4D and Fig. S5). Thus, our human GC models generated by the workflow faithfully reproduce within-cell type similarities as well as degree-of-pathology differences measured in our experiments.

The development of realistic conductance-based models and the inclusion of 15 distinct ionic conductances shown to express in human GC neurons offers a computational framework to study disease progression changes at a mechanistic level. Pairwise comparison between WG1 vs. WG4 models pointed to three conductances exhibiting the most prominent, WG-related difference: Ca-dependent K-channel (BK), Cav2.2 and Kir2.1. Specifically, we found that reducing BK- and increasing Cav2.2- and Kir2.1-conductance from their respective WG1 value (percentual change of median conductance from WG1 to WG4: BK, 56% increase; Cav2.2, 28% decrease; Kir2.1: 52% decrease) resulted in electrophysiology properties closely resembling WG4 cases (Fig. 4E, left; Fig. S6). To test these model predictions, we also performed single-cell RNA-sequencing in the resected hippocampal tissue of a subset of our patient cohort (4 out of 7 patients; 2 WG1, 2 WG4) and quantified the expression of genes associated with the three conductances, namely KCNMA1 (BK), CACNA1B (Cav2.2) and KCNJ2 (Kir2.1). We found that the trends (size effect) predicted by the models for the two ionic conductances are supported by the RNA-seq data (their absolute numbers expressed in unique molecular identifiers or UMI), i.e. we observed upregulation of BK and downregulation of Cav2.2 with disease progression (Fig. 4E). The direction of changes associated with Kir2.1 was inconclusive due to low number of gene reads.

How do the three identified ion conductances affect changes observed in electrophysiology properties, i.e. the f-I slope and time-to-first-spike? We used the aforementioned single-neuron models and performed a sensititivity analysis by perturbing parameters such as somatic Cav2.2 and Kir2.1 conductances and measuring their impact on the f-I slope and time-to-first-spike (Fig. S7). Importantly, the conductance perturbations in this analysis were similar to the ones brought about by the disease progression. Out of the three conductances, we found that only the decrease in somatic Cav2.2 from WG1 to WG4 affected the f-I gain in the same manner as observed experimentally (Fig. 2 and S7). Moreover, a decrease in dendritic Kir2.1 conductance was the only perturbation that resulted in a reduction of spike latency (Fig. S7). It follows that changes in somatic Cav2.2 and dendritic Kir2.1 conductances brought about by the disease best explain the changes observed in the most salient electrophysiology properties, respectively.

## Circuit excitability

How are the differences observed at the single-cell level manifested in a network setting? We used the single-neuron models to generate a network mimicking key features of DG circuitry, consisting of biophysically realistic and connected excitatory (GC) and inhibitory (human basket cells, BCs) neurons (network: 500 GC models, 6 human BCs (*18*)) (Fig. 5A). We tested the role of WG-dependent alterations on network dynamics via two independent manipulations: (i) by altering the network GC-composition between WG1 and WG4 GC models while maintaining all other aspects of the network identical (i.e. BC composition, connectivity, synaptic weight, external input, etc.), and (ii) by doubling the synaptic density between GCs from WG1 to WG4 (Fig. 5B). While Poisson-like, external input resulted in asynchronous GC and BC output for an unconnected network (“WG1 no syn”), instantiating recurrent connectivity between WG1 GCs and BCs (“WG1”) resulted in recurring, burst activity. We use burst activity as a proxy for hippocampal circuit excitability. Substituting WG1 GC models with WG4 ones while preserving all other aspects of the circuit (“WG4”) resulted in increased frequency of burst occurrence (Fig. 5B-C). Markedly, increased excitability for the “WG4” network occurred despite WG4 GCs exhibiting decreased spike-frequency gain (Fig. 2F and 4C) compared to WG1. This can be attributed to their shorter reaction time (i.e. broadly related to the ‘time-to-spike’ feature) which effectively translates into reduced rheobase. Following our observation of the approximate doubling of spine density from WG1 to WG4 (Fig. 3F), we instantiated a 2-fold increase in recurrent connectivity between WG4 GCs in the network (“WG4 x2”) that resulted in even stronger, sharper and more frequent burst events (Fig. 5B-C). Importantly, network excitability is substantially reversed when the three ionic conductances found to be differentially expressed between WG1 and WG4 GCs (BK, Cav2.2, Kir2.1) are substituted in “WG4” GC-models with their “WG1”-like values while all other aspects of the network remained unperturbed (“WG4 alt”). We conclude that GC-specific alterations observed during TLE-progression in the human hippocampus can readily lead to increased, recurring circuit excitability congruent with clinical observations of patients with increased HS. Moreover, combined, disease-related alterations in BK, Cav2.2 and Kir2.1 conductances predicted via data-driven, biophysical modeling and supported by independent, single-cell RNA-seq analysis, critically influence circuit dynamics dictating the transition from suppressed (WG1) to elevated (WG4) circuit excitability.

**Figure 5:**
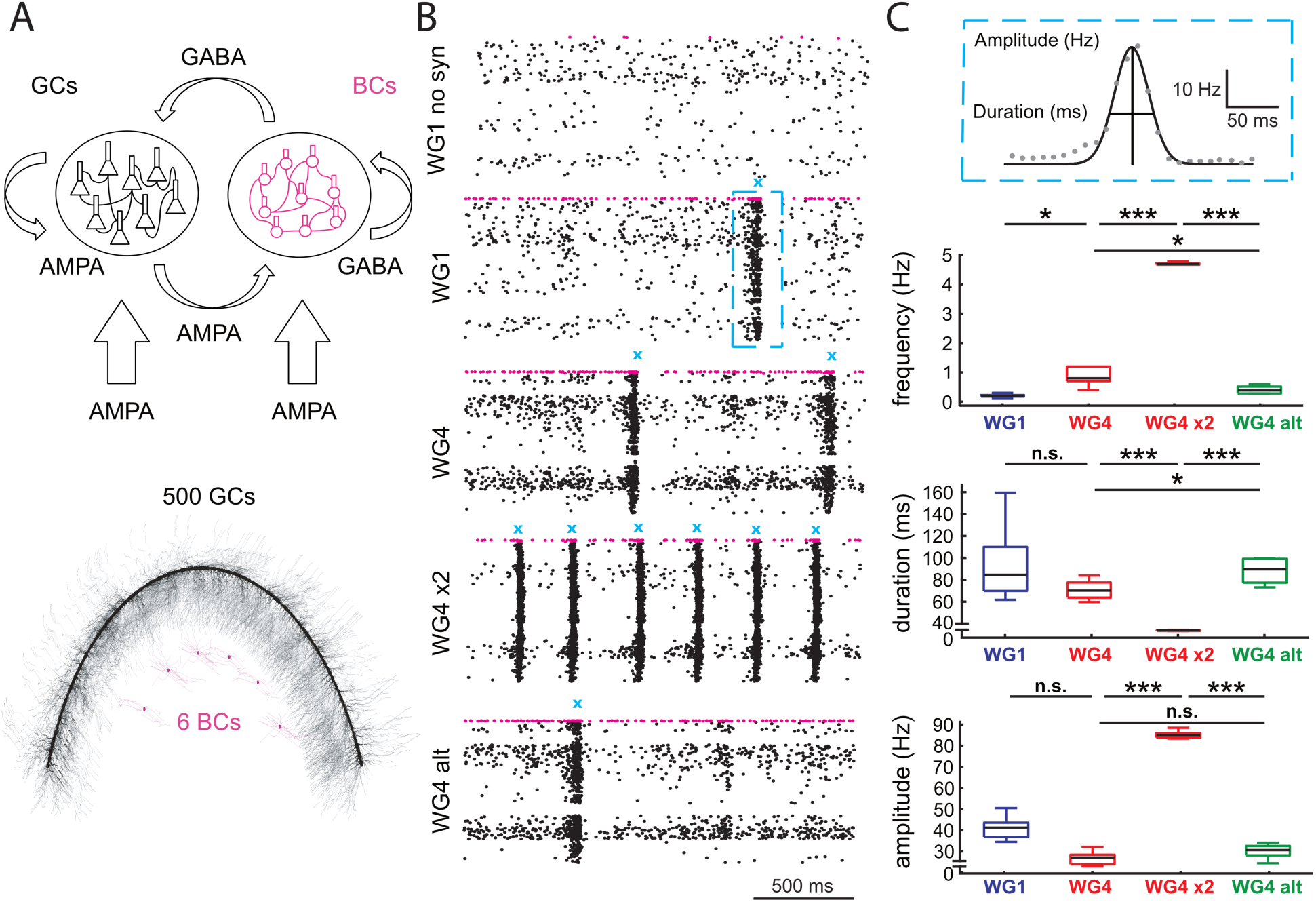
Systematic alterations in granule cell properties with disease progression reproduce increased pathophysiological excitability in a model of human dentate gyrus. (A) DG network model consisting of human single-cell models: 500 GCs (black) and 6 BCs (magenta). (Top) Arrows indicate the synaptic connectivity of excitatory (AMPA) and inhibitory (GABA) synapses. Large arrows on the bottom correspond to excitatory synaptic drive from the perforant pathway, smaller arrows signify within-network synaptic connectivity (see Materials and Methods). (Bottom) Morphologically and topographically realistic depiction of the DG network structure. (B) Raster plots of network dynamics during 2 s of activity driven by perforant path, Poisson-like input (total simulated time: 30 s; black: GC spikes, magenta: BC spikes). (Top to bottom) Unconnected network (only feedforward, “WG1 no syn”); WG1-type network with recurrent connectivity (“WG1”); WG4-type network identical to previous one (“WG1”) but with all WG1 GC models substituted with WG4 GC models (“WG4”); same network as before (“WG4”) but with double the recurrent connectivity between GCs (“WG4 x2”); same network as “WG4” but with altered set of ionic conductances (green in Figure 4A) setting them to their WG1-value (“WG4 alt”). Network dynamics in the presence of recurrent connectivity results in recurrent burst activity (BA; marked in cyan). (C) Analysis of BA event properties for various DG network configurations. In total, 5 network permutations, each with shuffled connectivity between GCs, were instantiated and simulated for each network configuration. (Top) Definition of three BA event properties: event frequency (i.e., number of BA-events per unit of simulated time), event amplitude and duration (as defined by fitting a Gaussian envelope to each event; grey: average GC firing rate measured in 10 ms time bins; black: Gaussian fit of BA-event envelope). (Below) BA feature analysis for WG-dependent network configuration (blue: “WG1”; red: “WG4” and “WG4 x2”; green: “WG4 alt”; line: mean; box plots: s.e.m. across 5 network realizations for each condition). Statistical testing via 1-way ANOVA with statistical significance: *: *p* < 0.05, ** *p* < 10^−3^; ***: *p* < 10^−4^.

## Discussion

How cells in the human brain change with disease progression and become hyperexcitable contributing to temporal lobe epilepsy seizures remains unanswered. This is particularly true for neurons in the human hippocampus, a brain region tightly linked to seizure initiation and support. We found that human granule cells, a prominent excitatory cell type in the hippocampus that survives disease progression, alter a number of their properties that are significantly affected by TLE and HS progression as witnessed in their genomic signature. How are these changes manifested? Electrophysiologically, the most prominent difference in GCs with disease progression is the decreased f-I gain. Yet, shorter spike latency of WG4 GCs vs. their WG1 counterparts leads to elevated, recurring circuit excitability (despite reduced f-I gain), an observation in-line with clinical observations in patients with increased HS typically suffering from increased number of seizures (*4–8*) and an earlier stage of onset (*8, 14, 15*). Our works suggests that alteration of GC and network excitability with disease progression is linked via single-cell RNA-seq and modeling to three ionic conductances, BK, Cav2.2, Kir2.1.

In the context of human epilepsies, BK channels have gathered considerable attention, e.g. (*28*). Yet, gain-of-function of BK exhibited in our patient cohort as well as described in patients of genetic epilepsies has been difficult to explain mechanistically (*29–31*). Here, we offer a mechanistic framework relating gain-of-function in BK channels of human granule cells, key regulators of hippocampal excitability (*20*), with increased network excitability. Our findings in human GCs also broadly agree with observations in rodents, in which a seizure-induced switch in BK-channels results in an excitability increase of cortical and DG neurons (*32–34*).

Concurrently, our single-cell perturbation analysis attributed changes in f-I slope and time-to-first-spike in the reduction of the Cav2.2 and Kir2.1 conductance, respectively, during TLE progression. Also referred to as N-type Ca current, it has been implicated in TLE with a number of anti-convulsant medications impacting it (*35–38*). Notably, our data suggest that, to revert disease progression, a gain-of-function in Cav2.2 is required. Alteration in the expression level of Kir2-channels has also been linked to excitability changes of granule cells in TLE (*39, 40*). While we detected WG-dependent Kir2-conductance differences in our GC models, RNA-seq revealed that overall expression of these channels is comparatively low (Fig. 4B and 4E; in agreement with (*41*)).

We observed significant differences in the morphology and spine density of GCs with disease progression with GCs from severely sclerotic cases exhibiting thicker dendrites and double the spine density (*42*). Despite clear indication that recurrent excitation between GCs impacts network excitability and dynamics, our simulations, in agreement with experiments in rodents (*43*), demonstrate that three ionic conductances, if reverted to their WG1-level (i.e. reducing the BK and increasing the Cav2.2 and Kir2.1 conductance by approx. 30-50%), can substantially reduce network excitability. This emerging mechanistic understanding of differences brought about with disease progression underlying TLE, points toward improved therapies such as viral delivery in a cell type-specific manner of genes for specific electrical conductances that can reverse the deleterious pathophysiology effects of epilepsy.

## Acknowledgements

We wish to thank Allen Institute founder, Paul G. Allen, for his vision, encouragement, and support. We thank Brandon Blanchard and Kael Dai for assisting with illustrations, Soo Yeun Lee, Tom Chartrand, Yina Wei, Bosiljka Tasic for discussions, and Lynne Becker for transitional project management support. We thank the Allen Institute Tissue Procurement team, especially Nick Dee and Julie Nyhus, the sequencing core, especially Kimberly Smith, Jeffy Goldy and Amy Torkelson, and Facilities team members for supporting human surgical tissue acquisition and transport. We thank the Allen Institute Histology team for processing of biocytin filled neurons and immunohistochemistry, and Imaging team for generating 20X slice scans and 63X Z-stack images for neuron reconstruction work, especially Samuel D Lee, Melissa Gorham, Kiet Ngo, Nadezhda Dotson, Lydia Potekhina, Nathalie Gaudreault, Medea McGraw and Rusty Nicovich. We thank Tim Jarsky and Luke Campagnola for development and support of the data acquisition software MIES. We are grateful to Caryl Tongco and Jae-Guen Yoon at Swedish Neuroscience Institute/Swedish Medical Center for coordinating patient consent, patient metadata, and tissue collections. We thank Dr. Carter Gerard for discussions and consultation early in the project, Dr. Ivan Soltesz for discussions and suggested analyses, and Drs. Rostad and Driscoll for providing de-identified patient tissue pathology reports.

## Author contributions

conceptualization: CAA, JT; data generation: AB, AN, RDF, PC, RM, BK, RG, SB, JT, RH; data curation: JB, CAA, JT, RDF, SMC, SS; formal analysis: AB, RDF and CAA; funding acquisition: CAA, JT, CK; investigation: AB, JT and CAA; project administration: CAA, JT, SMC, CK; writing original work: AB and CAA; writing – review and editing: all authors.

