## Supplemental material and methods for "Multi-modal characterization and simulation of human epileptic circuitry"

[x] Equal author contribution

† Current address: Institute for Advanced Clinical Trials for Children, WA, USA

### Supporting material and methods

#### Tissue preparation

Human brain tissue from neurosurgical origin was made available through the generosity of tissue donors (Table S1). Excised human brain tissue was collected from the brain within 1-3 minutes of resection (up to 5 minutes in rare cases), and transported in chilled, oxygenated ACSF.VII from the hospital to the Allen Institute laboratories within 15-30 minutes. Tissue was then mounted for slice preparation on the chuck of a Compresstome VF-200 or VF-300 vibrating microtome (Precisionary Instruments) to be sliced perpendicular to pial surface. Each human tissue slice was mounted in the recording chamber and inspected to ensure that the entire hippocampal depth was intact. Regions of the hippocampus were identified visually. Dentate gyrus granule cells along the granule cell layer were targeted for the full dataset. More details on tissue preparation can be found in (*43*) under section “Electrophysiology overview”.

#### Single nucleus RNA sequencing analysis

Intact nuclei from the hippocampus of four human tissue donors—H16.06.008 (WG1), H17.06.015 (WG1), H16.06.009 (WG4) and H16.06.010 (WG4) — were collected and processed for sequencing using 10X Genomics as follows. To generate single nuclei, flash frozen hippocampal samples were Dounce homogenized, stained with mouse anti-NeuN antibody (Millipore, FCMAB317PE), and NeuN+ (~70%) and NeuN- (~30%) nuclei were collected using fluorescence activated cell sorting (FACS). Sorted nuclei suspensions were concentrated to ~1000 nuclei/µl and loaded onto a 10X Genomics version 3 single-cell RNA-sequencing chip with a targeted capture of ~8000 nuclei per specimen. Nuclei (n = 29,212) were sequenced at a median read depth of 83,474 ± 24,111 reads/nucleus (median ± standard deviation). Median gene detection was 5295 ± 240 genes/nucleus and the median number of UMIs/nucleus was 16,088 ± 1790. For more information on RNA-seq technology applied in human brain tissue, see (*44*, *45*).

RNA-seq data was curated to remove nuclei with less than 1'000 genes detected, leaving 27,242 nuclei remaining. Nuclei were then assigned to broad classes (GCs, other excitatory, inhibitory, and non-neuronal) via supervised clustering, using the 100 most selective genes for each class. GC genes were defined as genes with the most significant differential expression between "DG" and "HiF" in the "Differential Search" in the Allen Human Brain Atlas (http://human.brain-map.org/ (*24*)). Similarly, markers for the other classes were selected using the RNA-seq Data Navigator: Human—a part of the Allen Cell Types Database (http://celltypes.brain-map.org (*45*)) — by running "Find Marker Genes" for "Cell Category" against all other cells. This identified genes specific for each class based on single nucleus RNA-seq in human middle temporal gyrus. The RNA-seq data was normalized by taking the unique molecular identifier (UMI) count for each cell, calculating counts per million (CPM), and then converting to logarithmic space: EXPR_norm = log2(CPM(UMI)+1). We then performed supervised *k*-means clustering with *k*=4 on all normalized data for nuclei using only these differential genes to identify cells most likely corresponding to each cell class. 12'011, 6'386, 5'302, and 3'543 nuclei were found corresponding to GCs, other excitatory, inhibitory, and non-neuronal cells, respectively. Expression of select ion channels across broad classes is shown in Figure 4. Following analysis of the computational models (see below), gene expression in GCs was analyzed for genes coding for the following ion channels: KCNMA1 (BK), CACNA1b (Cav2.2) and KCNJ2 (Kir 2.1). Box plots showing gene expression for each donor are presented in Figure 4 without a statistical assessment due to the small sample size for this type of analysis (N=4 donors).

**Electrophysiological recordings**

For electrophysiological recordings, human brain slices were mounted in a custom designed chamber and held in place with a slice anchor (Warner Instruments). The slice was bathed in ACSF at 34±1°C (warmed by an npi hpt-2 flow-through heater and thermafoil, controlled by an npi TC-20) at a rate of 2 ml per minute (Gilson Minipuls 3 pump). As a target temperature, 34°C was chosen to approximate physiological conditions, but with a safety buffer so as to not exceed 37°C. Bath temperature was continuously monitored. ACSF oxygenation was maintained by bubbling 95% O2, 5% CO2 gas in the specimen reservoir as well as delivering across the surface of the incubation bath via the custom designed specimen chamber. Solution oxygen levels, temperature and flow rate were monitored; values were measured, documented, and calibrated weekly to verify system quality.

Thick walled borosilicate glass (Sutter BF150-86-10) electrodes were manufactured (Sutter P1000 electrode puller) with a resistance of 3 to 7 MΩ. Prior to recording, electrodes were filled with 20 µl of internal solution SOP consisting of potassium gluconate, HEPES and other components (*43*) with biocytin, which was thawed fresh every day and kept on ice. The pipette was mounted on a Multiclamp 700B amplifier headstage (Molecular Devices) fixed to a micromanipulator (PatchStar, Scientifica).

Electrophysiology signals were recorded using an ITC-18 Data Acquisition Interface (HEKA). Commands were generated, signals processed, and amplifier metadata was acquired using a custom acquisition software program, written in Igor Pro (Wavemetrics). Data were filtered

(Bessel) at 10 kHz and digitized at 50 kHz. Data were reported uncorrected for the measured -14 mV liquid junction potential between the electrode and bath solutions.

Upon break-in and formation of a stable seal (typically within the first minute and not more than 3 minutes after break-in), the resting membrane potential of the neuron was recorded. All recordings were bridge balanced and systematically checked for access resistance matching the quality control criteria described in (*46*) under section "Electrophysiology overview".

**Morphology reconstruction and feature extraction**

Morphological reconstructions were generated using previously described methods (*43*) under section "Morphology and Histology Overview". Briefly, biocytin-filled neurons were stained via diaminobenzidine reaction and imaged at 63x magnification on a Zeiss Axio Imager 2. Individual cells were digitally reconstructed with custom-written software (Vaa3D and Mozak) to create accurate, whole-neuron representations saved in the SWC format. A feature extraction suite was adopted and customized enabling the analysis of dendritic and somatic reconstructions and extraction of a number of features related to branching pattern, size, density, soma position, etc. resulting in 59 morphology features (Table S4).

Dendritic spine densities were assessed using Neurolucida 360 (v2017.01.1) and Imaris (v9.3). Initial and terminal 100 µm sections of the dendritic region outside the granule cell layer were visualized for manual spine density estimation along 10 µm-long segments per dendritic branch.

### Data analysis

We used the standard Python libraries for analyzing the electrophysiological and morphological data: Mann-Whitney U-testing (mannwhitneyu from statistics library), random forest classification (RandomForestClassifier with 200 decision trees from scikit-learn lirary) and regression analysis (linalg.lstsq from numpy). For dimensionality reduction to perform tSNE we used the sklearn.manifold package.

### Electrophysiology feature extraction and single-cell model setup

From whole-cell patch-clamping experiments, electrophysiological responses to a battery of standardized current stimuli (1 s-long dc current injections of increasing amplitude) were analyzed resulting in a set of subthreshold and spiking features for each experiment (Table S2). Electrophysiology features such as spike timing, amplitude, width, etc. were obtained for each experiment. For a subset of features, regression was used to assess how a particular electrophysiology feature changes with increasing intracellular stimulation amplitude. In total, 30 electrophysiology features were extracted from our *in vitro* experiments (Table S2) and analyzed with a number of statistical techniques for WG-dependent differences. Details on the implementation of the feature extraction analysis and relevant code is publicly available through (*47*).

The model optimization procedure for generating biophysically realistic single-neuron models is feature-based and attempts to set the somatic, axon initial segment, and dendritic properties in such manner to capture features of intracellular somatic responses for a number of standardized current stimulation waveforms. For model generation, electrophysiology features were extracted using a feature extraction library (eFEL) developed in (*48*). Specifically, 11 electrophysiology features were extracted for each experiment (Fig. S4) and their mean and standard deviation (std) was computed for a particular stimulation waveform. If more than one sweep of the same stimulation waveform exists (majority of the experiments), then the std of that particular waveform was used. If only a single sweep of the experiment exists, a default value of 10% was used for the std.

The compartments of human granule and basket cells were separated into three zones: the axon initial segment (AIS), the soma and dendrites. In our models, the AIS was represented by one fixed-length section with a total length of 30 μm and 1 μm diameter.

In all models, passive and active properties were optimized using the same fitting procedure. For passive properties, one value for the specific capacitance (*cm*), passive conductance (*g_pas*), passive reversal potential (*e_pas*), and cytoplasmic resistivity (*Ra*) was uniformly set across all compartments. Notably, the values of these parameters were part of the genetic optimization procedure (see below). Active channel mechanisms were spatially uniformly distributed in the AIS, soma, and dendrites with every zone receiving a separate set of channels (Table S5). The same approach was used for generating biophysically detailed basket cell models from human medial temporal gyrus with slice electrophysiology and morphology data originating from (*46*) (specimen IDs: 529807751, 541536216; Table S6). The final excitatory and inhibitory models will be made available on https://github.com/AllenInstitute/epilepsy_human_dg.

**Parameter optimization and single-cell model generation workflow**

For the single-cell model optimization and generation, the BluePyOpt framework was adopted (https://github.com/AllenInstitute/All-active-Workflow, https://github.com/AllenInstitute/All-active-Manuscript) (*48*) which utilizes multi-objective optimization relying on an evolutionary optimization algorithm. Briefly, for every electrophysiological feature out of the 11 used to build computational models (Fig. S4), an absolute standard score was calculated $Z_i=|f_i-\mu_i|/\sigma_i$ with the feature value ($f_i$) measured from the output traces of the models and $\mu_i$, $\sigma_i$ being the experimentally measured mean and standard deviation, respectively (*48*).

The development of every biophysically detailed single-cell model involved a 3-stage optimization workflow that iteratively focused on a set of conductances. In the first stage, passive model properties like membrane resistance and capacitance were set by fitting subthrehold responses and subthreshold features such as *voltage_sag*, *voltage_base* and *voltage_after_stim*. In the second stage, 15 active ionic conductances were included while keeping the membrane capacitance and passive reversal potential fixed. Notably, the initial parameter ranges for all GC models (both for passive and active ion channels) were chosen irrespective of WG (Table S5) and optimized for 100 generations. In this stage, ten spiking traces were used for model training along with subthreshold responses. In the final stage, the best model from the previous stage was chosen, its parameter range extended by two-fold for all conductances and optimization was re-initialized for at least 50 generations. As part of the BluePyOpt framework, multiple models were generated for every cell with the best individual considered as the one with the smallest sum of objective values. The best model for each experiment was used in the pool of models selected for the network simulations (see below). Concurrently, for every cell, the 10 best models ("hall of fame" models according to (*25*, *48*)) were selected and used for pairwise conductance comparisons between WG1 and WG4 (Fig. S6).

For the evolutionary algorithm, a population size of typically 320 individuals on 320 cores of a BlueGene/P or Cori Haswell (Cray XC40) supercomputer was used. Generation of a single, biophysically detailed, single-neuron model required approximately 200'000 core-hours of computational time on the aforementioned architectures.

**Hippocampal network model setup**

A bio-plausible DG network model was constructed consisting of 506 single-cell models with the details of the network relying on (*18*). Briefly, GC and BC cells in our network were positioned via a topographic approach with somata of the models positioned in 3D space along a ring with a radius of 1500 μm for GCs and 750 μm for BCs. The ring was opened to form the arch roughly corresponding to the physical extent of human DG (Fig. 5A).

Synaptic connectivity in the network depends on the location of a neuron along the ring (characterized by the arch-angle). The overall neural network structure was adapted from (*18*). We have modified the number of connections per synapse to make it more realistic and in agreement to (*49*). The following connectivity structure was chosen to represent the DG circuit with periodic boundary conditions (compensating for border artifacts). Every GC formed 50 connections with its 100 closest GCs along the ring with the exact connectivity between the postsynaptic target pool chosen randomly. Every GC also projected to 3 of its closest BCs along the ring. Recurrent GC-GC connections formed 2 to 5 synapses with the exact number chosen randomly. The location of GC synapses was chosen along dendrites within 150 μm from the soma. All excitatory connections of GCs corresponded to AMPA synapse kinetics (*18*).

Every BC created 100 out of 140 possible connections with its topographically closest GCs with synapses located on the soma of GCs. Recurrent BC connectivity was also implemented so that every BC is connected to its two closest neighbors with the exact location of inhibitory synapses chosen randomly within 150 μm from the BC soma. All inhibitory connections correspond to GABA-A synapse kinetics (*18*). Synaptic kinetics were approximated using biexponential synapse models (`exp2syn' in (*50*)) involving two time constants ($\tau_1$ and $\tau_2$) as well as the reversal potential ($V_{rev}$). The values used for these parameters are the following (adopted from (*18*)):

| *synapse type* | $\tau_1$ / *ms* | $\tau_2$ / *ms* | $V_{rev}$ / *mV* |
|---|---|---|---|
| GC → GC (AMPA) | 1.5 | 1.5 | 0.0 |
| GC → BC (AMPA) | 0.3 | 0.6 | 0.0 |
| Perforant path → GC | 1.5 | 5.5 | 0.0 |
| BC → BC (GABA-A) | 0.16 | 1.8 | -70.0 |
| BC → GC (GABA-A) | 0.26 | 5.5 | -70.0 |
| Perforant path → BC | 2.0 | 6.3 | 0.0 |

To activate the network, GCs and BCs received background synaptic input along their dendrites. GCs received 5 external connections per cell with 5 to 15 synapses each – the exact number of synapses was chosen randomly from a homogenous distribution. BCs received the same amount of background synaptic input. Every background connection was activated by a Poisson process (rate: 3 Hz) emulating perforant path input. Background synaptic input corresponded to AMPA synapse kinetics both for GCs and BCs (*18*).

All network simulations were setup and performed using inhouse developed software based on (*51*). The versions of the DG network presented in Fig. 5 will be made publicly available through https://github.com/AllenInstitute/epilepsy_human_dg

**Supplementary figure 1**

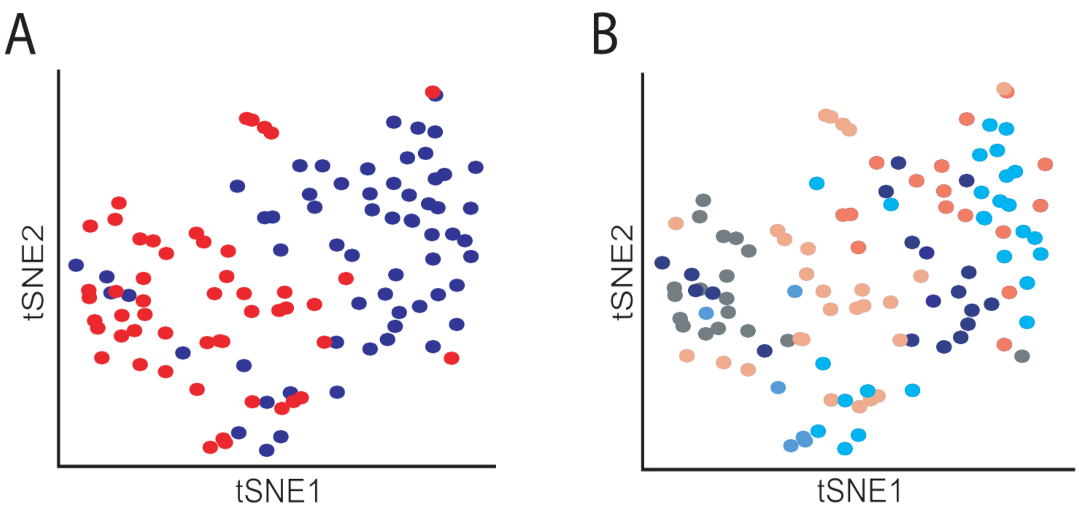


**Supplementary figure 1: Patient-specific electrophysiological properties.** (A) tSNE visualization of 30 electrophysiology features per cell for 112 DG GCs from all subjects (blue: WG1; red: WG4; 4 WG1, 3 WG4 subjects; Table S1). (B) Same as panel (A) with colors corresponding to different patients.

**Supplementary figure 2**

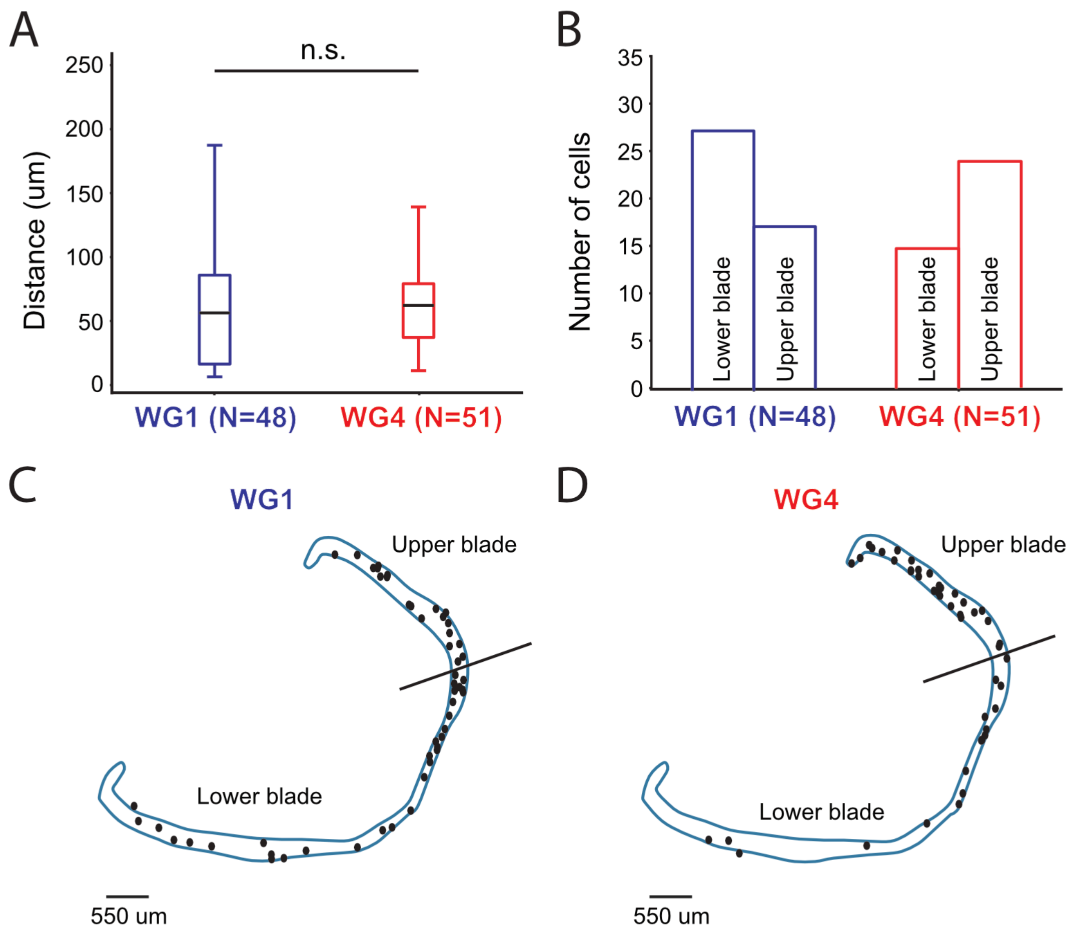


**Supplementary figure 2: Location of sampled granule cells within human dentate gyrus.** (A) Distance between the soma of whole-cell patch-clamped and morphologically reconstructed DGs from the inner side of DG. Statistical significance testing (Mann-Whitney U-test, significance level = 0.05) shows no difference in soma depth location of sampled neurons across DG. (B) Number of sampled GCs located in the upper vs. lower DG blade indicate the absence of a location-bias. Data are from all reconstructed cells (102) for the entire patient cohort (7 patients, Table S1). (C), (D) Raw data for the depth and soma location (upper/lower blade) of sampled GCs within the GC-layer for WG1 and WG4. The separating line indicates the fictitious landmark separating upper and lower blade in our analysis.

**Supplementary figure 3**

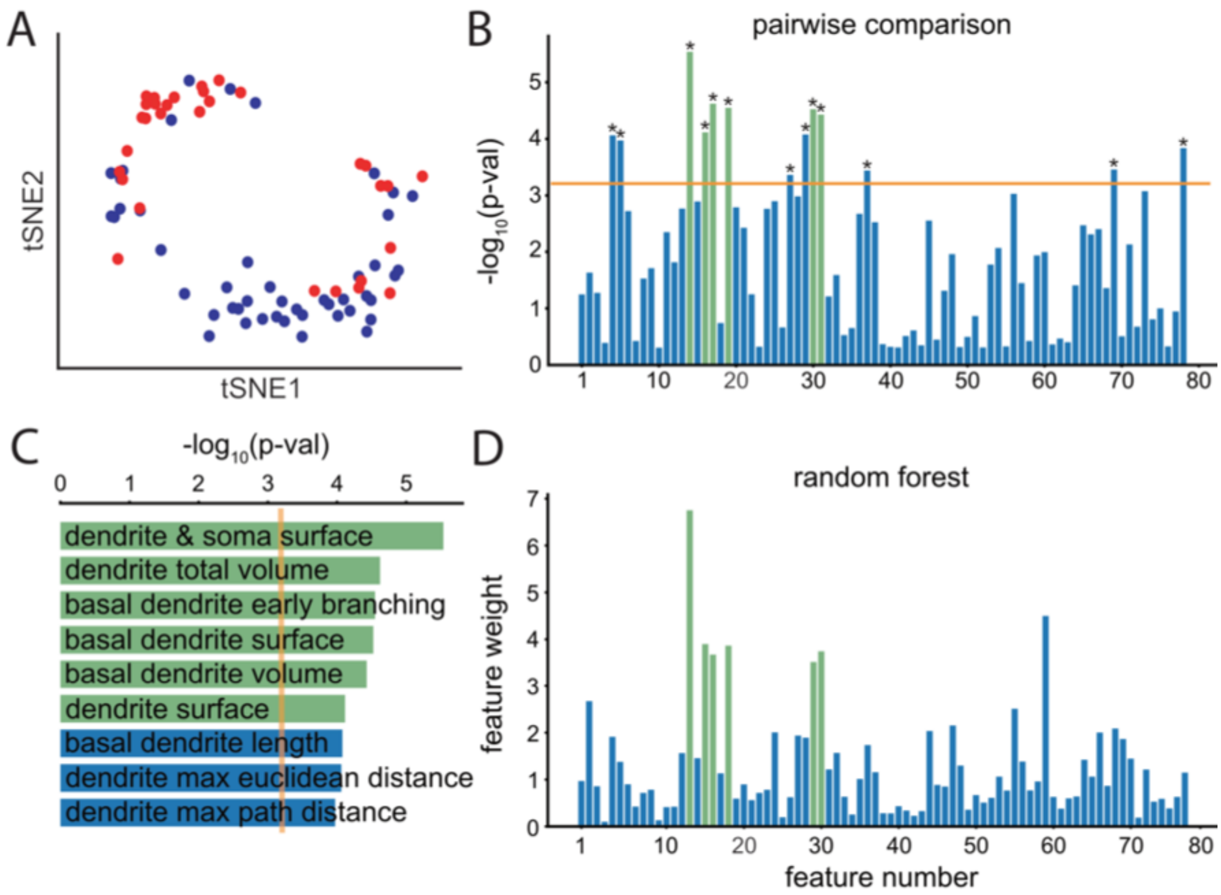


**Supplementary figure 3. Classification of both electrophysiological and morphological features with degree of sclerosis.** (A) tSNE visualization of electrophysiological and morphological features (blue: WG1; red: WG4) of GCs with both data modalities present (77 GCs out of 112 electrophysiologically recorded cells and 102 morphologically reconstructed had both data modalities, i.e. 68% out of all possible electrophysiology experiments and 75% out of all morphology reconstructions). (B) Pairwise comparison of WG1 and WG4 between 79 morpho-electric features (orange line: significance level $p$-value = 0.05 / 79 = 0.0006; stars indicate features of statistical signifance). (C) 9 most significant morpho-electric features between WG1 and WG4 cells (order determined per the *p*-value from the pairwise comparisons in panel B). (D) Feature weights of random forest classifier trained on the same set of morpho-electric features as in panel B (green boxes: features shared between pairwise comparison, panel B, and the random forest analysis, panel D).

**Supplementary figure 4**

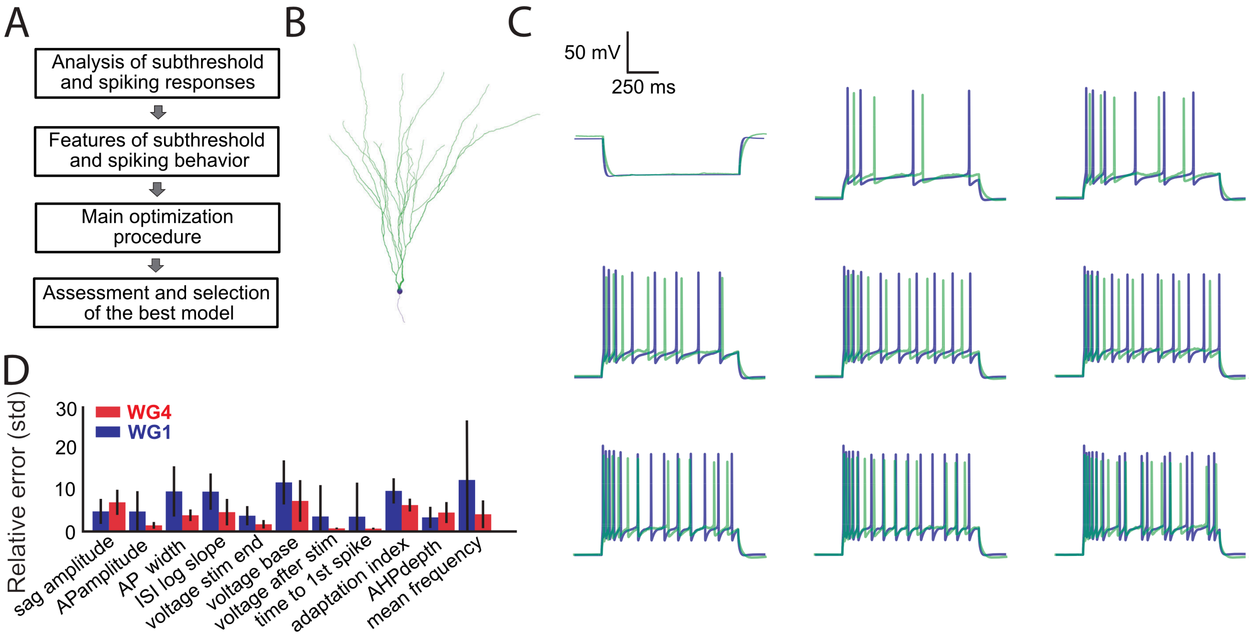


**Supplementary figure 4. Development of the biophysically detailed single-neuron models from slice electrophysiology data and morphology reconstructions.** (A) Individual steps of the main optimization procedure. (B) Reconstructed single-neuron morphology of a human DG granule cell. (C) The granule cell from the same experiment (panel B) is subjected to 1s-long current injections and its electrophysiological responses measured via whole cell patch-clamping. Comparison between experimental electrophysiological traces (blue) and the best biophysical GC model (green) generated by the 3-step optimization procedure from the particular experiment (Materials and Methods). (D) Average model error for each of the 11 electrophysiological features chosen for generation of the single-cell models averaged over all available trials. The 3-stage optimization workflow is based on BluePyOpt (*48*). The relative error is measured in number of standard deviations from the experimental traces (blue: WG1 models; red: WG4 models). In the boxplots, the bars correspond to mean values and black lines designate the standard deviation of the relative error across all WG1 and WG4 single-cell models.

### Supplementary figure 5

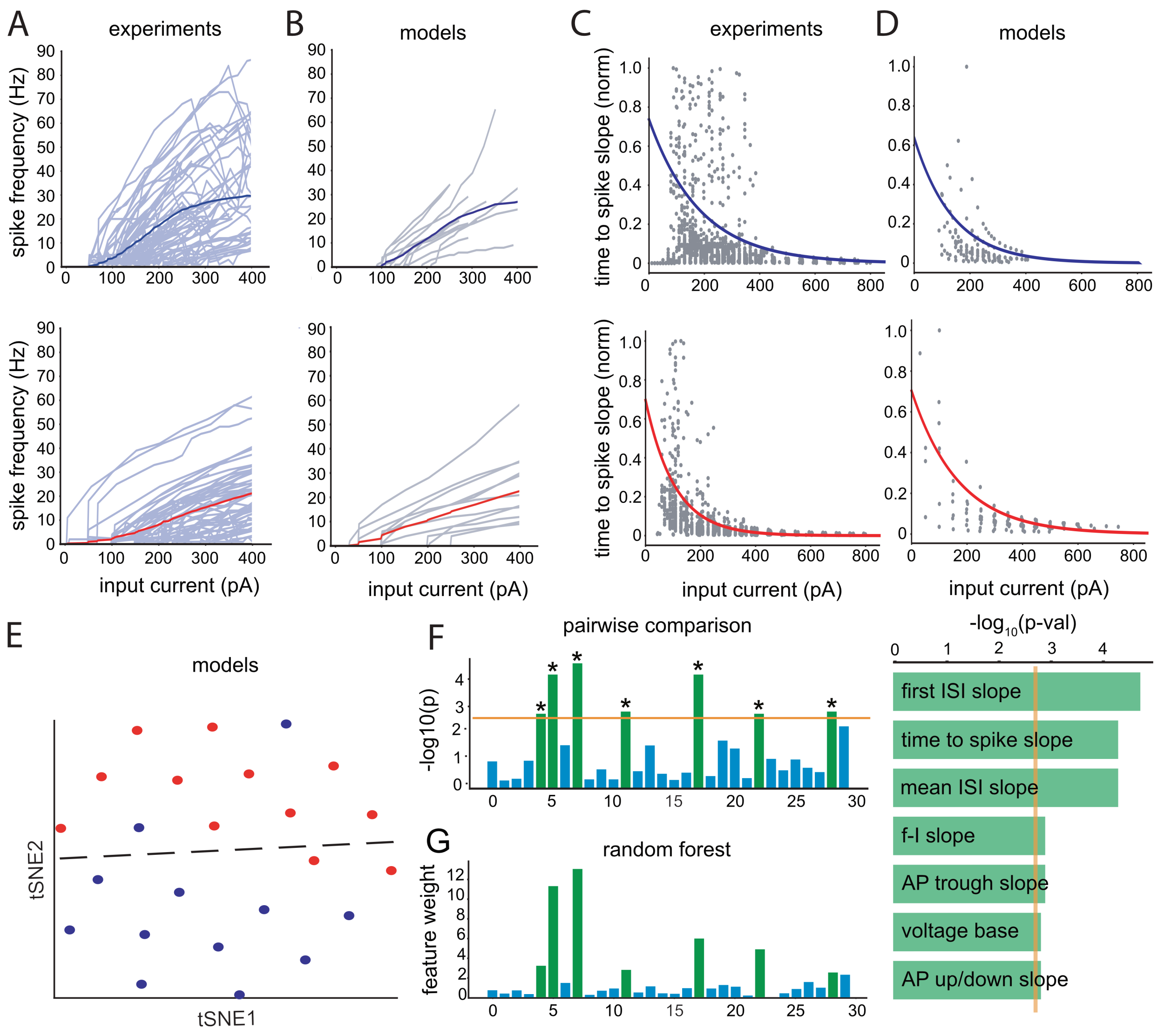


**Supplementary figure 5: Side-by-side comparison of electrophysiology features between experiments and single-cell models with disease progression.** Spike frequency and time-to-spike as response to increasing somatic injection current for mild (WG1; blue) vs. severe (WG4; red) degree of sclerosis. (A) f-I data from whole-cell patch-clamp experiments (*n*=112 human GCs in total, WG1: 61 GCs, WG4: 51 GCs; grey: individual human GC experiments; thick line:

mean). (B) f-I data from single-cell GC model simulations (*n*=24 human GC models in total, WG1: 12 GC models, WG4: 12 GC models; grey: responses from individual GC models developed from experiments; thick line: mean). (C) Normalized time-to-spike as response to increasing suprathreshold somatic current injection from whole-cell patch-clamp experiments (circles: time-to-spike in a particular sweep; thick line: exponential fit, $\tau_{exp}$(WG1)=0.005±0.0003, $\tau_{exp}$(WG4)=0.009±0.0002). (D) Normalized time-to-spike responses from single-cell GC model simulations (*n*=24 human GC models in total, WG1: 12 GC models, WG4: 12 GC models; grey: responses from individual GC models developed from experiments; thick line: exponential fit, $\tau_{model}$(WG1)=0.007±0.001; $\tau_{model}$(WG4)=0.006±0.0006). (E) tSNE visualization of the electrophysiology features from the GC models based on their response to 1 s-long somatic dc current injections (blue circles: WG1, red: WG4; left, broken line: *k*-means clustering). Clear separation between WG1 vs. WG4 GC-models is observed based on their electrophysiology responses as is the case for experiments (Fig. 2, B and C). (F) Pairwise comparison of electrophysiological features between WG1 and WG4 GC-models across feature space (left) and in decreasing level of statistical significance (right). Statistical significance is calculated using Mann-Whitney U-test with the significance level (orange line) Bonferroni-corrected for the number of features (* designates $p$-value < 0.05/30=0.002). (G) Electrophysiology feature weights of random forest classifier trained on the simulated electrophysiology features. As observed, electrophysiology features between pairwise comparison and random forest classifier are shared. Importantly, the four most significant features leading the separation between WG1 and WG4 GC-models (first ISI slope, time-to-spike slope, mean ISI slope and f-I slope) are the same ones separating the experimental data (compare with Fig. 2, D and E). It follows that the GC-models developed through the 3-stage optimization workflow (Fig. S4) capture the most prominent differences between WG1 and WG4 GCs as measured in our experiments.

**Supplementary figure 6**

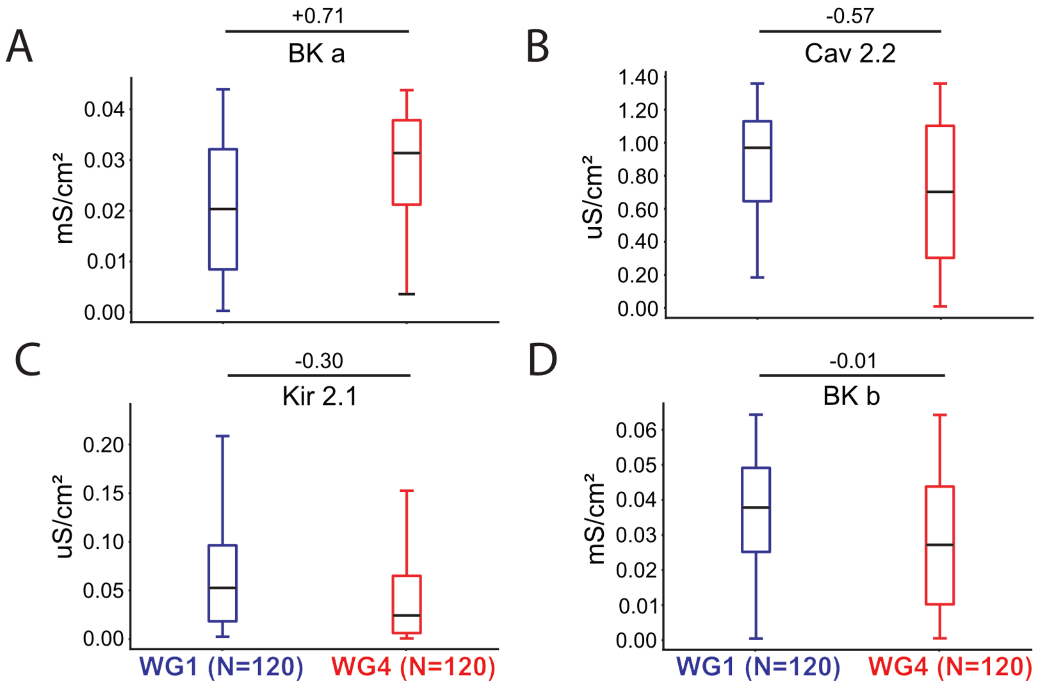


**Supplementary figure 6: Comparison of ionic conductance densities between WG1 and WG4 single-neuron models.** For a particular GC experiment, the 3-stage-optimization workflow is used and the 10 best models ("hall of fame" models according to (*48*)) are selected (instead of only the best) and used for pairwise conductance comparisons between WG1 and WG4. Notably, the BK conductance used in our biophysical models is separated into two parts, one without the b4 accessory subunit (BKa) exhibiting slower kinetics and one with it (BKb)(*23*, *52*). Pairwise comparison between WG1 and WG4 granule cell models (120 WG1, 120 WG4 models) reveals statistically significant differences between four conductances: (A) somatic BKa, (B) somatic Cav2.2, (C) dendritic Kir2.1, and (D) axonal BKb. Statistical significance differences between WG1 and WG4 models are assessed via Mann-Whitney U-testing, Bonferroni-corrected for multiple comparisons (45 comparisons per model for active conductances across morphology zones). The size effect (Cohen's *d*) is shown for the four conductances that exhibit the most prominent differences. The fact that these conductances are critically implicated in alterations of neural dynamics with disease progression is further established when reverting them from their WG4- to their WG1-values reduces network excitability (Fig. 5).

**Supplementary figure 7**

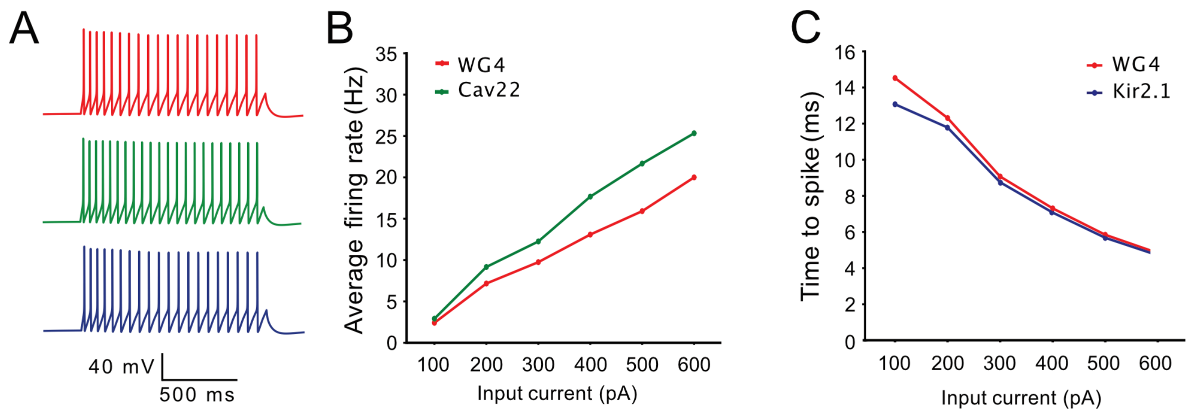


**Supplementary figure 7: Sensitivity analysis via biophysical modeling linking specific, conductance-based perturbations to electrophysiological characteristics.** (A) Somatic voltage traces of individual models in response to intracellular current injection. The same single-cell model of WG4 (developed as part of Fig. 4, main manuscript) is shown unperturbed (red), with altered somatic Cav2.2 conductance (green) and altered dendritic Kir2.1 (blue). (B) Focus on the impact of the somatic Cav 2.2 conductance on f-I slope. Action potential response to intracellular current dc injections (duration: 1 s) of increasing amplitude (circles: mean frequency response of 12 models). Red – WG4 models as developed for Figure 4 (main manuscript). Green – WG4 models with increased somatic Cav 2.2 conductance by 56% mimicking the conductance-value of WG1 models (compare to Fig. S6B). Increase in the Ca-conductance results in the reduction of the f-I slope. (C) Focus on the impact of the dendritic Kir2.1 conductance on time-to-first-spike (spike latency). Spike latency during the current step stimulation for the same models as in B (circles: mean frequency response of 12 models). Red – original WG4 models. Green - WG4 models with increased Kir 2.1 conductance by 52% compared to WG4 (compare to Fig. S6C). Increase in Kir-conductance results in decrease of the spike latency for weak current injection amplitude with the effect disappearing for larger injection amplitudes.

**Table S1**

| Patient ID | Age | Sex | Seizure onset (age) | IQ | Hemisphere | Seizure rate (per month) | Seizure type | Clinical history | WG | # cells with electrophysiology recordings | # cells with morphology reconstructions |
|---|---|---|---|---|---|---|---|---|---|---|---|
| H16.06.008 | 24 | F | 17 | | Left | | Partial, Generalized | | 1 | 2 | 2 |
| H16.06.013 | 34 | F | 20 | | Left | | Partial | | 4 | 6 | 14 |
| H17.06.012 | 23 | M | 20 | | Right | | Partial | | 3 | 18 | 15 |
| H17.06.014 | 44 | M | 1-3 | 79 | Left | 1-2 | Partial | Tuberous sclerosis, Traumatic brain injury | 1 | 16 | 14 |
| H17.06.015 | 19 | M | 12 | 89 | | 3-5 | Partial, Generalized | | 1 | 17 | 15 |
| H18.06.366 | 38 | M | childhood | | Left | 1-2 | Partial, Generalized | Developmental delay, ~7 y. o. | 4 | 27 | 21 |
| H18.06.368 | 59 | M | 17 and 57 | 101 | Left | 1-2 | Partial, Generalized | Multiple seizure types | 1 | 26 | 24 |

**Table S1.** Patient metadata and experimental yield.

**Table S2**

| Feature # | Electrophysiology feature names | Electrophysiology feature description |
|---|---|---|
| 1 | adaptation_slope | adaptation index, current slope (1/pA) |
| 2 | adaptation_rheobase | adaptation index, rheobase current (pA) |
| 3 | AP_trough_rheobase | AP trrought, rheobase current (pA) |
| 4 | AP_downstroke_slope | AP downstroke, current slope (dV/dt/pA) |
| 5 | voltage_base_mean | voltage baseline (mV) |
| 6 | time_to_spike_slope | time to first spike, current slope (ms) |
| 7 | R_in | input resistance (MOhm) |
| 8 | first_ISI_slope | first ISI, current slope (ms/pA) |
| 9 | AP_thr_rheobase | first AP threshold, rheobase current (pA) |
| 10 | AP_upstroke_slope | first AP upstroke, current slope (dV/dt/pA) |
| 11 | voltage_base_sigma | voltage std (mV) |
| 12 | AP_trough_slope | first AP through, current slope (mV/pA) |
| 13 | time_to_spike_rheobase | time to first spike, current slope (pA) |
| 14 | AP_height_slope | first AP height, current slope (mV/pA) |
| 15 | AP_downstroke_rheobase | first AP downstroke, current rheobase (pA) |
| 16 | AP_up/downstroke_rheobase | AP up/downstroke, rheobase current (1/pA) |
| 17 | taum_mean | membrane time constant (ms) |
| 18 | mean_ISI_slope | mean ISI, current slope (ms/pA) |
| 19 | AP_width_slope | AP width, current slope (ms/pA) |
| 20 | AP_thr_slope | AP threshold, current slope (mV/pA) |
| 21 | first_ISI _rheobase | first ISI, current rheobase (pA) |
| 22 | AP_height_rheobase | first AP height, rheobase current (pA) |
| 23 | AP_up/downstroke_slope | first AP up/downstroke, current slope (1/pA) |
| 24 | AP_width_rheobase | first AP width, rheobase current (pA) |
| 25 | mean_ISI_srheobase | mean ISI, current rheobase (pA) |
| 26 | AP_upstroke_rheobase | first AP upstroke, current rheobase (pA) |

| | | |
|---|---|---|
| 27 | taum_sigma | membrane time constant std (ms) |
| 28 | rheobase_current | rheobase current (pA) |
| 29 | fI_slope | f-I curve slope (Hz/pA) |
| 30 | rheobase_freq | AP frequency at rheobase (Hz) |
| 31 | voltage_sag | Averaged voltage sag (mV) |

**Table S2.** In vitro electrophysiology features extracted from the experimental whole-cell patch-clamp experiments. A short description of each feature is offered while a more detailed one can be found in (*43*).

**Table S3**

| PATIENT CASE | CLASSIFIER PREDICTION | NUMBER OF CELLS | PREDICTED WG | TRUE WG |
|---|---|---|---|---|
| H16.06.008 | 0.50 | 2 | WG1 | WG1 |
| H16.06.013 | 1.00 | 6 | WG4 | WG4 |
| H17.06.012 | 0.94 | 18 | WG1 | WG4 |
| H17.06.014 | 0.68 | 16 | WG1 | WG1 |
| H17.06.015 | 0.58 | 17 | WG1 | WG1 |
| H18.06.366 | 0.77 | 27 | WG1 | WG4 |
| H18.06.368 | 0.76 | 26 | WG1 | WG1 |

**Table S3.** Patient-out-validation of the electrophysiology features based on WG. Random forest classifier trained on data from 6 patients aiming to predict WG-score of the 7$^{th}$ patient based on cellular electrophysiology features (Table S2). Ground-truth WG-score was determined via expert pathologist scoring. Classifier prediction refers to the fraction of cells attributed the predicted WG-score by the classifier. The WG-score which > 50% of cells of a case are ascribed to constitutes the predicted WG-score for that case.

**Table S4**

| Feat # | Feature description | Feat # | Feature description |
|---|---|---|---|
| **1** | dendrite_average_diameter | **31** | basal_dendrite_total_surface |
| **2** | dendrite_contraction | **32** | basal_dendrite_total_volume |
| **3** | dendrite_early_branch | **33** | apical_dendrite_average_diameter |
| **4** | dendrite_max_branch_order | **34** | apical_dendrite_contraction |
| **5** | dendrite_max_euclidean_distance | **35** | apical_dendrite_early_branch |
| **6** | dendrite_max_path_distance | **36** | apical_dendrite_max_branch_order |
| **7** | dendrite_mean_parent_daughter_ratio | **37** | apical_dendrite_max_euclidean_distance |
| **8** | dendrite_neurites_over_branches | **38** | apical_dendrite_max_path_distance |
| **9** | dendrite_num_bifurcations | **39** | apical_dendrite_mean_parent_daughter_ratio |
| **10** | dendrite_num_branches | **40** | apical_dendrite_neurites_over_branches |
| **11** | dendrite_num_outer_bifurcations | **41** | apical_dendrite_num_bifurcations |
| **12** | dendrite_num_stems | **42** | apical_dendrite_num_branches |
| **13** | dendrite_num_tips | **43** | apical_dendrite_num_outer_bifurcations |
| **14** | dendrite_parent_daughter_ratio | **44** | apical_dendrite_num_stems |
| **15** | dendrite_soma_surface | **45** | apical_dendrite_num_tips |
| **16** | dendrite_total_length | **46** | apical_dendrite_parent_daughter_ratio |
| **17** | dendrite_total_surface | **47** | apical_dendrite_total_length |
| **18** | dendrite_total_volume | **48** | apical_dendrite_total_surface |
| **19** | basal_dendrite_average_diameter | **49** | apical_dendrite_total_volume |
| **20** | basal_dendrite_early_branch | | |
| **21** | basal_dendrite_max_branch_order | | |
| **22** | basal_dendrite_max_euclidean_distance | | |
| **23** | basal_dendrite_max_path_distance | | |
| **24** | basal_dendrite_neurites_over_branches | | |
| **25** | basal_dendrite_num_bifurcations | | |
| **26** | basal_dendrite_num_branches | | |
| **27** | basal_dendrite_num_outer_bifurcations | | |
| **28** | basal_dendrite_num_stems | | |
| **29** | basal_dendrite_num_tips | | |
| **30** | basal_dendrite_total_length | | |

**Table S4.** Single-cell morphological features extracted from granule cells used to classify between WG1 and WG4. The table provides the names of all features. For a detailed description of each feature, see (*43*).

**Table S5**

| | AIS | Soma | Dendrites |
|---|---|---|---|
| **Membrane capacitance** | cm (0.1 : 5 μF/cm$^2$) | | |
| **Membrane resistance (passive)** | g_pas (1e-7 : 1e-2 S/cm$^2$) | | |
| **Passive reversal potential** | e_pas (-110 : -70 mV) | | |
| **Axial resistance** | Ra (50 : 1000 Ω-cm) | | |
| **Potassium Kir2.1** (Kir21_gc.mod) | gkbar_Kir21 (1e-7 : 1e-2 S/cm$^2$) | | gkbar_Kir21 (1e-7 : 1e-2 S/cm$^2$) |
| **Potassium Kv4.2** (Kv42_gc.mod) | | | gkbar_Kir21 (1e-7 : 1e-2 S/cm$^2$) |
| **Calcium Cav1.2** (Cav12_gc.mod) | gbar_Cav12 (1e-7 : 1e-2 S/cm$^2$) | gbar_Cav12 (1e-7 : 1e-2 S/cm$^2$) | gbar_Cav12 (1e-7 : 1e-2 S/cm$^2$) |
| **Calcium Cav1.3** (Cav13_gc.mod) | gbar_Cav13 (1e-7 : 1e-2 S/cm$^2$) | gbar_Cav13 (1e-7 : 1e-2 S/cm$^2$) | gbar_Cav13 (1e-7 : 1e-2 S/cm$^2$) |
| **Calcium Cav2.2** (Cav22_gc.mod) | gbar_Cav22 (1e-7 : 1e-2 S/cm$^2$) | gbar_Cav22 (1e-7 : 1e-2 S/cm$^2$) | gbar_Cav22 (1e-7 : 1e-2 S/cm$^2$) |
| **Calcium Cav3.2** (Cav32_gc.mod) | gbar_Cav32 (1e-7 : 1e-2 S/cm$^2$) | gbar_Cav32 (1e-7 : 1e-2 S/cm$^2$) | gbar_Cav32 (1e-7 : 1e-2 S/cm$^2$) |
| **gkbar_SK2** (SK2_gc.mod) | gkbar_SK2 (1e-7 : 1e-2 S/cm$^2$) | | gkbar_SK2 (1e-7 : 1e-2 S/cm$^2$) |
| **HCN (Ih)** (HCN_gc.mod) | | | gbar_HCN (1e-7 : 1e-2 S/cm$^2$) |
| **Calcium buffer** (Cabuffer_gc.mod) | tau_Cabuffer (20 : 1000 ms) | tau_Cabuffer (20 : 1000 ms) | tau_Cabuffer (Cabuffer.mod) (20 : 1000 ms) |
| **Ca buffer** (Cabuffer_gc.mod) | brat_Cabuffer (100 : 1000 ms) | brat_Cabuffer (100 : 1000 ms) | brat_Cabuffer (100 : 1000) |
| **Potassium Kv11** (Kv11_gc.mod) | gkbar_Kv11 (1e-7 : 1e-2 S/cm$^2$) | | |
| **Potassium Kv14** (Kv14_gc.mod) | gkbar_Kv14 (1e-7 : 1e-2 S/cm$^2$) | | |
| **Potassium Kv34** (Kv34_gc.mod) | gkbar_Kv34 (1e-7 : 1e-2 S/cm$^2$) | | |
| **Potassium Kv723** (Kv723_gc.mod) | gkbar_ Kv723 (1e-7 : 1e-2 S/cm$^2$) | | |
| **Ca-dependent potassium BK** (BK_gc.mod) | gakbar_BK (1e-7 : 1e-2 S/cm$^2$) | gakbar_BK (1e-7 : 1e-2 S/cm$^2$) | |
| **Potassium BK** (BK_gc.mod) | gabkbar_BK (1e-7 : 1e-2 S/cm$^2$) | gabkbar_BK (1e-7 : 1e-2 S/cm$^2$) | |
| **Ca-dependept potassium SK2** (BK_gc.mod) | gkbar_SK2 (1e-7 : 1e-2 S/cm$^2$) | | gkbar_SK2 (1e-7 : 1e-2 S/cm$^2$) |
| **Sodium Na8st** (na8st_gc.mod) | gbar_na8st (1e-7 : 1e-2 S/cm$^2$) | gbar_na8st (1e-7 : 1e-2 S/cm$^2$) | |
| **Potassium Kv21** (Kv21_gc.mod) | | gbar_na8st (1e-7 : 1e-2 S/cm$^2$) | |

**Table S5.** Ionic conductance parameter ranges used to initialize model optimization of biophysically realistic, human granule cell models. Parameter ranges are reported along three morphology sections (AIS, soma and dendrites). Notably, the same parameter ranges were used to initialize the optimization workflow for all GC models irrespective of WG.

**Table S6**

| | AIS | Soma | Dendrites |
|---|---|---|---|
| **Membrane capacitance** | cm (0.5 : 10 μF/$cm^2$) | | |
| **Membrane resistance (passive)** | g_pas (1e-7 : 1e-2 S/$cm^2$) | | |
| **Passive reversal potential** | se_pas (-110 : -60 mV) | | |
| **Axial resistance** | Ra (50 : 150 Ω-cm) | | |
| **Sodium** (NaV.mod) | gbar_NaV (1e-7 : 5e-2 S/$cm^2$) | gbar_NaV (1e-7 : 5e-2 S/$cm^2$) | gbar_NaV (1e-7 : 5e-2 S/$cm^2$) |
| **Transient potassium** (K_T.mod) | gbar_K_T (1e-7 : 1e-2 S/$cm^2$) | | |
| **Potassium delayed rectifier** (Kd.mod) | gbar_Kd (1e-7 : 1e-2 S/$cm^2$) | | |
| **Potassium Kv2-like** (Kv2like.mod) | gbar_Kv2like (1e-7 : 1e-1 S/$cm^2$) | | |
| **Potassium Kv3-like** (Kv3_1.mod) | gbar_Kv3_1 (1e-7 : 1 S/$cm^2$) | gbar_Kv3_1 (1e-7 : 1 S/$cm^2$) | gbar_Kv3_1 (1e-7 : 1 S/$cm^2$) |
| **Potassium SK-type** (SK.mod) | gbar_SK (1e-7 : 1e-2 S/$cm^2$) | gbar_SK (1e-7 : 1e-2 S/$cm^2$) | |
| **Low-voltage activated calcium** (Ca_LVA.mod) | gbar_Ca_LVA (1e-7 : 1e-2 S/$cm^2$) | gbar_Ca_LVA (1e-7 : 1e-2 S/$cm^2$) | |
| **High-voltage activated calcium** (Ca_HVA.mod) | gbar_Ca_HVA (1e-7 : 1e-4 S/$cm^2$) | gbar_Ca_HVA (1e-7 : 1e-4 S/$cm^2$) | |
| **Calcium dynamics** (CaDynamics.mod) | gamma_CaDynamics (5e-4 : 5e-2 %) | gamma_CaDynamics (5e-4 : 5e-2 %) | |
| | decay_CaDynamics (20 : 1000 ms) | decay_CaDynamics (20 : 1000 ms) | |
| **HCN (Ih)** (Ih.mod) | | gbar_Ih (1e-7 : 1e-5 S/$cm^2$) | gbar_Ih (1e-7 : 1e-5 S/$cm^2$) |
| **Potassium m-like** (Im_v2.mod) | | | gbar_Im_v2 (1e-7 : 1e-2 S/$cm^2$) |

**Table S6.** Ionic conductance parameter ranges used to initialize model optimization of biophysically realistic, human basket models (BCs). Parameter ranges are reported along three morphology sections (AIS, soma and dendrites).
